# INTEGRATOR:Structural Elucidation of the INO80 Chromatin Remodeler via Experimentally Guided Molecular Simulations

**DOI:** 10.64898/2026.05.15.725493

**Authors:** Jules Nde, Gihan Panapitiya, Margaret S. Cheung, C Mark Maupin, Mihaela E. Sardiu

**Affiliations:** Department of Physics, University of Washington, Seattle, WA, 98195, USA; Applied Materials & Manufacturing Group, Pacific Northwest National Laboratory, Richland, WA, 99354, USA; Environmental Molecular Sciences Laboratory, Pacific Northwest National Laboratory, Richland, WA, 99352, USA; Chemical and Biological Signatures Group, Pacific Northwest National Laboratory, Richland, WA, 99352, USA; Department of Biostatistics and Data Science, University of Kansas Medical Center, Kansas City, KS, 66103, USA

**Keywords:** Chromatin remodelers, INO80 complex, Modularity, integrated modeling

## Abstract

The INO80 chromatin remodeling complex plays a central role in DNA repair, transcription, and replication. Yet, a comprehensive understanding of its structural organization remains incomplete due to the dynamic nature of several of its subunits and the sharing of several subunits with related remodeling complexes. Here, we report a computational model of the three-dimensional structure of the *S. cerevisiae* INO80 complex using an integrative approach that combines experimental crosslinking mass spectrometry, molecular docking, and molecular dynamics simulations. Our results reveal the spatial and dynamical organization of key modules—ARP8, ARP5, NHP10, and RVB1/2—within the intact complex. The resulting structural model agrees with crosslinking constraints, highlighting the architecture of the previously uncharacterized NHP10 module. This module, including the C-terminal region of the Ino80 scaffolding protein, has remained elusive due to its intrinsic flexibility and lack of high-resolution structural data. To facilitate this integrative modeling workflow and make it broadly accessible, we presented INTEGRATOR: (INTEGRAtive TempOral and stRuctural Analysis of protein modules), a versatile workflow package designed as a tool to elucidate the structure and dynamics of large, flexible macromolecular assemblies using well-established softwares. Our findings demonstrate the power of integrative modeling in resolving the role of the highly disordered NPH10 module in recruiting other dynamic modules into INO80 large protein assemblies and offer a generalizable framework for determining the architecture of similarly complex and heterogeneous molecular machines. This work carries broad implications for understanding the structural basis of chromatin regulation in microbial organisms and the implications for the dysregulation in diseases such as cancer.

## 1 Introduction

Chromatin remodeling complexes (remodelers) are essential ATP-dependent molecular machines that control the structure and accessibility of chromatin during critical cellular processes such as transcription, DNA replication, and repair (1, 2). Among these, the INO80 complex is especially notable for its roles in repositioning nucleosomes, regulating histone variant exchange, and maintaining genome stability, thereby influencing gene expression and protecting against genomic instability (3). These mechanisms, which contribute to genomic instability, are a hallmark of cancer development(4). Studies have shown increased sensitivity to genotoxic stresses and defects in replication fork restart in INO80-deficient cells, linking the complex to tumor suppression roles (5). INO80 achieves these functions by binding to promoter regions and sliding nucleosomes, often preferring hexasomes over canonical nucleosomes, and is required for proper chromosome segregation and restoration of chromatin structure after DNA damage (6, 7). INO80’s structural complexity arises from its modular assembly of over 15 subunits, some shared with other chromatin remodeling complexes like SWR-C and RSC (8–10). This subunit overlap, particularly of Rvb1/2 ATPases, actin, and actin-related proteins (e.g. Arp4, Arp5, Arp8), creates substantial analytical challenges, as these shared elements can adopt distinct conformations and functions depending on their molecular context. Discerning INO80’s unique architecture, therefore, requires not just high-resolution structural methods but also comparative and orthogonal biochemical analyses. Many INO80 modules, including flexible N-terminal regions and accessory subunits, exhibit pronounced conformational heterogeneity and dynamic assembly, often complicated further by partial occupancy or subunit exchange with related complexes. Such factors make the assignment of subunit topology and mechanistic roles difficult based solely on static structures, underscoring the need for integrative structural approaches.

Structurally, INO80 is among the largest and most elaborate chromatin remodelers, with its core based around a Rvb1/Rvb2 AAA+ ATPase heterohexamer that acts as both assembly scaffold and stator (11, 12). Recent cryo-electron microscopy (cryo-EM) studies have elucidated the global arrangement of the evolutionarily conserved core, revealing multivalent interactions with both histone and DNA and defining key domains such as the Swi2/Snf2 ATPase motor, Arp5, Ies6, and various grappler elements involved in remodeling and substrate specificity (10, 13–18). These studies also uncovered differential recognition of histone variants and suggested unified mechanisms for nucleosome sliding, histone editing, and chromatin boundary setting (15).

Nevertheless, despite these advances, a high-resolution understanding of INO80’s complete structure and dynamic assembly remains elusive. Several modules and subunits, including the NHP10 module and the C-terminal region of the scaffold protein Ino80, have eluded structural characterization due to their pronounced conformational flexibility, transient interactions, and partial disorder (14, 18). This dynamic nature hampers conventional crystallography, and even high-resolution cryo-EM has struggled to resolve these regions, resulting in persistent gaps in understanding how module architecture interrelates with enzymatic function.

Here, to overcome these obstacles, we present a comprehensive structural and mechanistic analysis of the yeast INO80 complex using an integrative approach that combines orthogonal experimental data (such as crosslinking mass spectrometry) with computational modeling and molecular dynamics. This strategy has emerged as powerful tools for building coherent models of flexible, and heterogeneous assemblies (19–23) for exploiting spatial restraints from crosslinking and global outlines from cryo-EM, but also employing in silico docking and dynamic simulations to capture alternative conformations and transient inter-subunit contacts. Despite its promising application, there are still technical gaps to solve the structure of a large complex with multiple modularities that vary with versatile remodeling purposes, such as INO80.

Our work incorporates crosslinking data, molecular docking, and atomistic simulation to resolve the spatial and dynamical organization of key modules—the ARP8, ARP5, NHP10, and RVB1/2 units—within the context of the intact complex. In particular, we define the previously elusive structure of the NHP10 module, gaining new insights into its association with the Ino80 scaffold and its potential regulatory functions. Our findings advance the field by bridging missing structural details and providing a dynamic, holistic 3D model of INO80, demonstrating the broad utility of integrative methods for complex, dynamic assemblies with core and peripheral modules. This framework (Figure 1) provides a robust platform for elucidating the mechanistic roles of chromatin remodelers in genome regulation and identifies potential avenues for understanding their dysregulation in human disease. Furthermore, we have encapsulated this integrative modeling workflow into a versatile software package named INTEGRATOR: (INTEGRAtive TempOral and stRuctural Analysis of pROtein modules), designed to facilitate the structural elucidation of large, dynamic macromolecular assemblies with multiple constituent modules.

**FIGURE 1.**
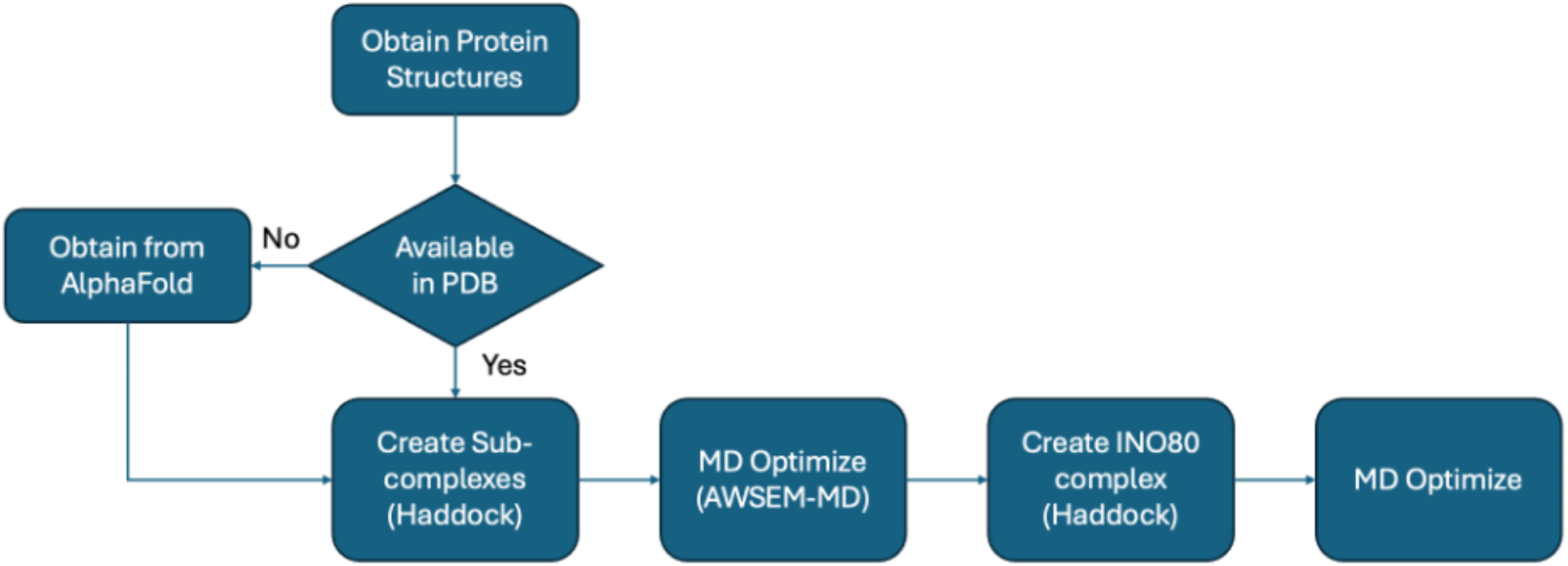
The workflow for predicting the INO80 structure involves several steps. First, the protein PDBs are obtained. If they are not available in the Protein Data Bank, they are retrieved from AlphaFold (24). Next, the complexes are created using HADDOCK, and MD optimization is performed on each sub-complex. Subsequently, HADDOCK is used to assemble the INO80 complex. Finally, MD optimization is conducted on the INO80 complex.

## 2 RESULTS

### 2.1 Overall structural organization of the INO80 complex

The INO80 chromatin remodeler is a large, multi-subunit complex in *Saccharomyces cerevisiae*, comprising over 15 unique subunits and reaching a combined molecular mass of approximately 1.3 MDa (Figure 2A). Of these, a core set of eight subunits is evolutionarily conserved from yeast to humans (Figure 2B), positioning INO80 as an important model for dissecting not only the assembly mechanisms of such megadalton complexes but also the structural underpinnings of conserved chromatin regulatory functions (Figure 2C-D) (25–28).

**FIGURE 2.**
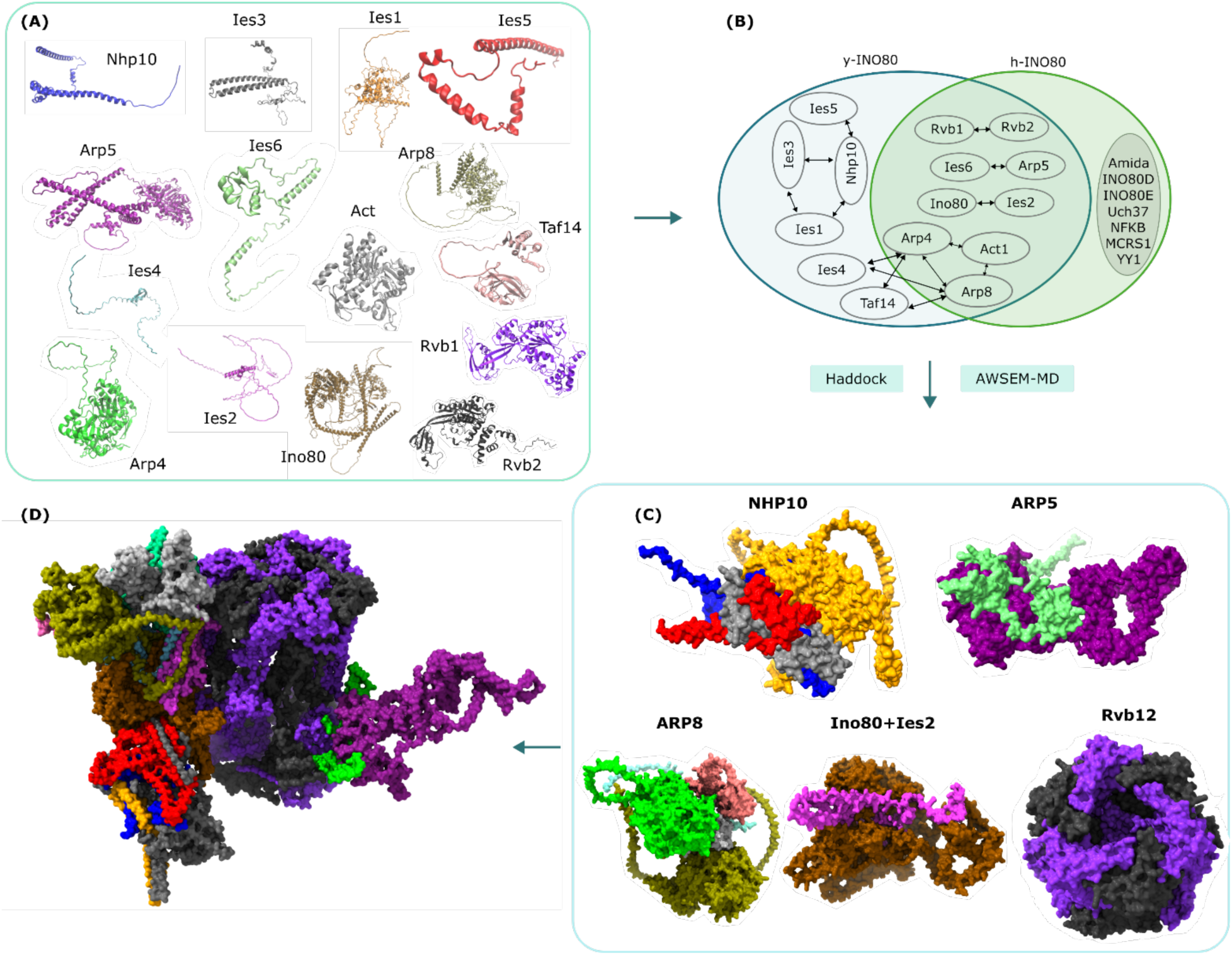
Workflow of the integrative modeling of the INO80 complex. (A) PDBs of the individual subunit of the INO80 complex (B). Shared and non-shared subunits of the yeast and human INO80 complex; (C) Individual modules of the INO80 complex (C) are assembled into the final complex (D). In the structural representation, the NHP10 module is depicted with Nhp10 in blue, Ies3 in red, Ies1 in grey, and Ies5 in orange. The ARP8 module is shown with Arp8 in tan, actin in silver, Arp4 in green, Taf14 in pink, and Ies4 in cyan. The ARP5 module includes Arp5 in purple and Ies6 in lime. Core components and additional subunits are color-coded as follows: Ino80 in ochre, Ies2 in magenta, Rvb1 in black, and Rvb2 in violet. This color scheme highlights the modular architecture of the INO80 complex and facilitates visual distinction between conserved modules and subunits across structural figures.

Within INO80 complex, the subunits are organized into four major functional modules—NHP10 (Nhp10, Ies1, Ies3, and Ies5), ARP8 (Arp8, Arp4, Act, Taf14 and Ies4), ARP5(Arp5, Ies6), and RVB1/2 (Rvb1, Rvb2) each grouped around a specific scaffolding protein that mediates the incorporation of accessory factors during complex assembly (10, 29). Deletion studies and biochemical analyses have established Ino80 protein as the central scaffold, coordinating the spatial arrangement and stability of all four subcomplexes, and linking ATPase activity to nucleosome remodeling and DNA repair functions (1, 30).

While structural and functional aspects of the ARP8, ARP5, and RVB1/2 modules have become increasingly well-characterized, the NHP10 module has posed a persistent challenge due to its high intrinsic flexibility, dynamic interactions, and limited structural data. Indeed, genetic and single-molecule studies have implicated the NHP10 module in DNA binding and the regulation of nucleosome sliding and length sensing, making its structural elucidation critical for understanding how INO80 achieves substrate specificity and processivity.

To address this knowledge gap, we applied an integrative structural modeling approach to resolve the spatial organization of INO80’s submodules. By leveraging crosslinking mass spectrometry, molecular docking, and molecular dynamics simulations, we provide new insights into the molecular architecture of the elusive NHP10 module and its integration with the other subcomplexes. Our results illuminate not only the modular assembly pathway of INO80 but also the distinct and conserved roles of each module in chromatin remodeling.

### 2.2 Assembly of the subcomplexes

#### 2.2.1 Structural Characterization of the NHP10 module

The NHP10 subcomplex comprises four unique subunits: Nhp10, Ies1, Ies3, and Ies5, with their individual structures predicted using AlphaFold2 and illustrated in Figure 3A. To construct an initial model of the NHP10 subcomplex, we applied molecular docking via HADDOCK(31), assembling the predicted individual subunit structures. Because the recruitment order and assembly pathway can influence the final complex conformation, two complementary strategies can be considered: experimental deletion studies to pinpoint the order of subunit incorporation, or extended molecular dynamics simulations to allow the initial model to relax and explore conformational space toward a stable structure.

**FIGURE 3.**
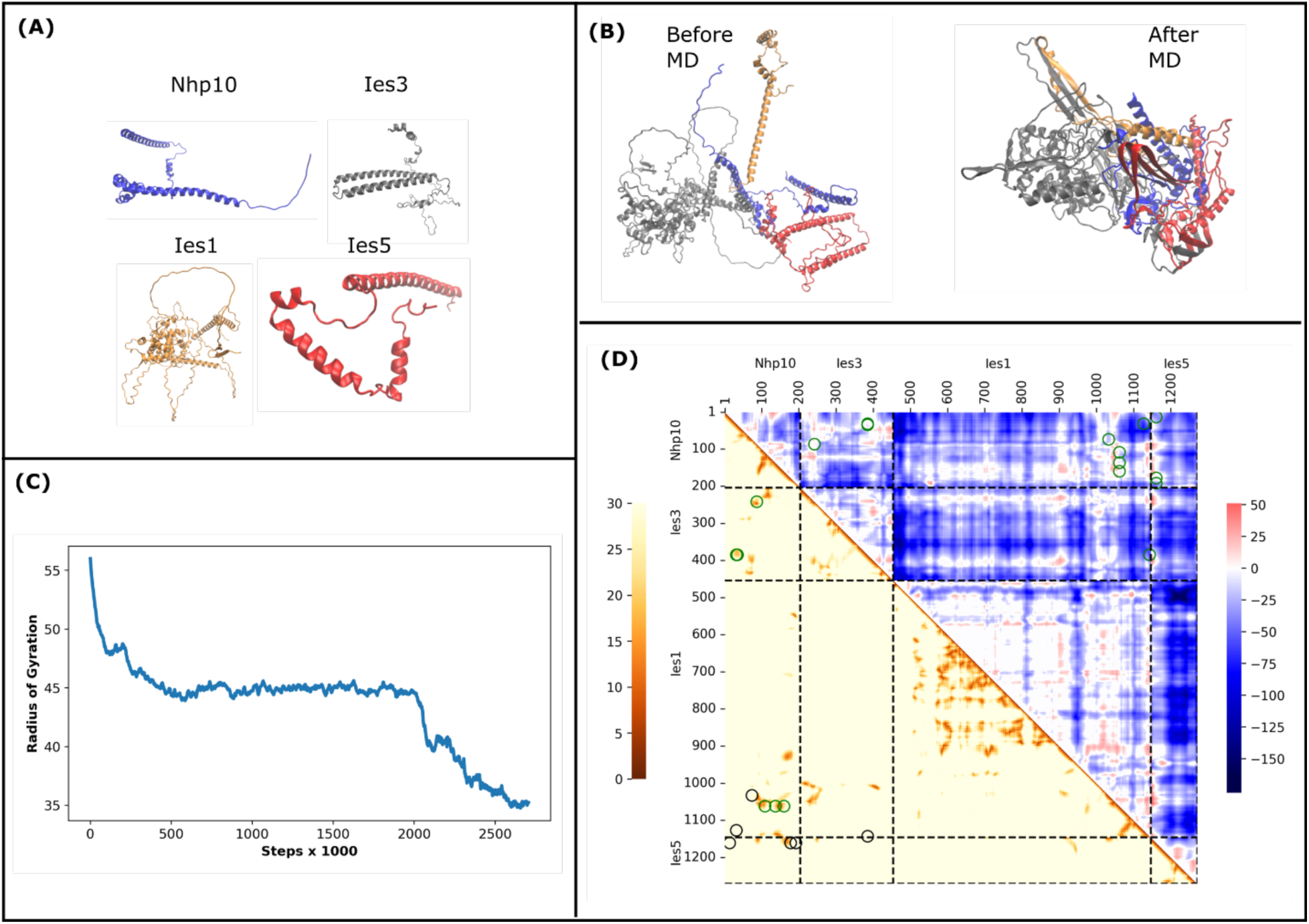
Workflow for Determining the NHP10 Module Structure: (A) Structures of the subunits of the ARP8 subcomplex are retrieved from AlphaFold2. (B) The individual structures are assembled to form the initial template using molecular docking. (C) The radius of gyration plot corresponding to the MD optimization. (D) Contact maps before and after MD optimization. The lower triangle shows the residue-residue distances; distances corresponding to the cross-links given in Tosi et al.(10) are marked in circles. Distances greater than 24 Å are colored in black, while others are in green. The upper triangle depicts the difference between final and initial cross-link distances. Negative differences indicate decreased distances, meaning residues moved closer together. Positive differences signify increased distances, indicating residues moved farther apart. For clarity, changes smaller than 10 Å have been set to zero. Both color bars represent values using Ångströms as the fundamental unit of measurement.

The Nhp10 subunit serves as the scaffold within this module, engaging in distinct physical interactions with Ies1, Ies3, and Ies5 (Figure 3B), consistent with prior biochemical studies. The NHP10 subcomplex is recruited into the larger INO80 assembly through interactions between the N-terminal domain of the Ino80 subunit and the C-terminal regions of the Ies1 and Ies3 subunits (Figure 3B and D). Crosslinking mass spectrometry additionally reveals contacts between the N-terminal regions of Ies3 and Ies2, suggesting an intricate network stabilizing subcomplex integration. Within the subcomplex, Ies5 associates peripherally with Nhp10, as confirmed by contact map analyses shown in Figure 3B and 3D. Notably, coarse-grained molecular dynamics simulations reveal the emergence of new contact interfaces not apparent in the initial docking model (Figure 3D). These contacts, corroborated by independent crosslinking data, underscore the significance of dynamic refinement to capture physiologically relevant interactions. The relaxation of the model during simulations is further supported by a decreased radius of gyration (Figure 3C), indicating a transition toward a more compact and stable conformation. This dynamic assembly likely contributes to the module’s regulatory capacity within INO80, potentially modulating chromatin interactions and remodeling activity in response to cellular contexts or DNA damage signals.

Together, these findings highlight the critical role of integrative modeling, combined with dynamic simulations, in resolving the flexible yet functionally essential regions of the INO80 complex’s architecture.

#### 2.2.2 Structural Characterization of the ARP8 module

Figure 4 illustrates the detailed predicted three-dimensional structure of the ARP8 module, a critical subcomplex within the large INO80 chromatin remodeling complex. Our structural analysis elucidates the overall organization of this module (Figure 4A), highlighting regions of pronounced flexibility enriched with key residues that facilitate protein-protein interactions, as visualized in the contact maps (Figure 4D). These interaction hotspots are corroborated by crosslinking mass spectrometry data, confirming their functional significance in stabilizing inter-subunit associations. To refine the structural model, we employed molecular dynamics simulations, which enabled relaxation of the module toward a more physiologically relevant conformation (Figure 4B). This relaxation was accompanied by an increase in inter-subunit contacts—as demonstrated by the difference in contact maps (Figure 4D)—and a reduction in the radius of gyration (Figure 4C), indicative of a more compact and stable assembly. Notably, many of the newly formed contacts observed in the MD-refined structure were independently validated by crosslinking experiments, underscoring the power of integrating computational modeling with experimental data. By integrating predicted structures with molecular dynamics simulations and experimental crosslinking data, we uncover how the ARP8 module acts as a critical sensor of extranucleosomal DNA, positioning near the entry site of the nucleosome and influencing substrate engagement. Our findings elucidate the pivotal role of ARP8 in modulating INO80’s nucleosome remodeling activity through conformational plasticity and module-specific interactions—insights that were previously elusive due to the module’s intrinsic flexibility and limited experimental resolution. The largely disordered appearance of Ies4 suggests a role in flexible inter-subunit tethering or as a regulatory hub facilitating conformational adaptation. The observed shifts in Arp8 and Act indicate how these subunits may undergo conformational adjustments, facilitating nucleosome recognition and energy transduction. Such flexibility is crucial as INO80 must remodel distinct chromatin contexts, including partially unwrapped or variant-containing nucleosomes, without dissociating. The distinct modular blocks corresponding to the Arp8, Act, Arp4, Taf14, and Ies4 subunits reveal tightly knit intra-module contacts essential for their stability and cooperative function. More importantly, inter-module contacts highlighted by off-diagonal clusters may represent allosteric interaction sites that propagate conformational changes spanning the complex, enabling coordinated ATPase regulation, DNA translocation, and histone-handling activities. Recent studies underline that communication between these modules is critical for INO80’s ability to mobilize nucleosomes efficiently while responding to various cellular signals and DNA damage cues.

**FIGURE 4.**
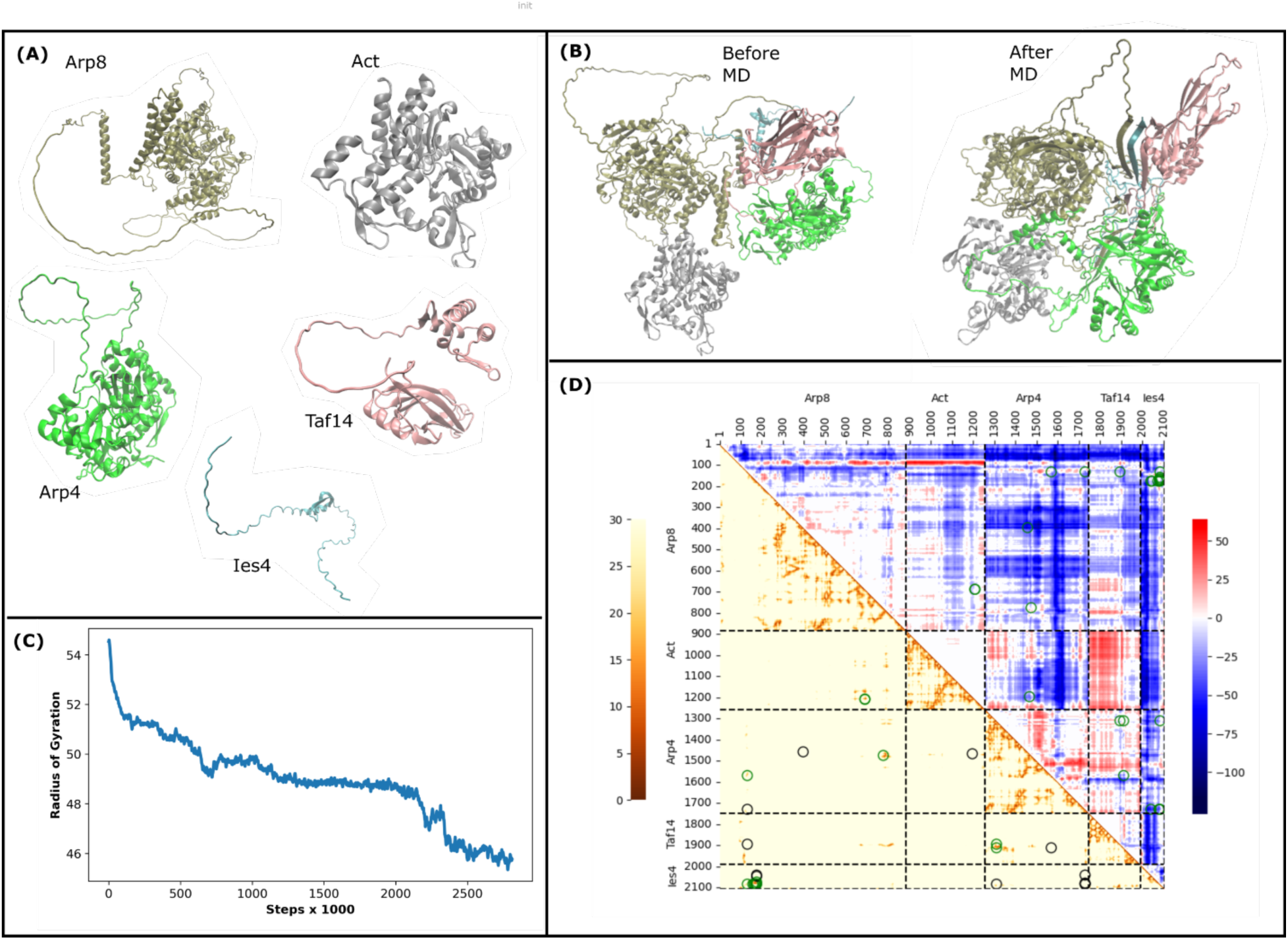
Workflow for Determining the ARP8 Module Structure: (A) Structures of the individual subunits of the ARP8 subcomplex are retrieved from AlphaFold2. (B) The individual structures are assembled to form the initial template using molecular docking. (C) The radius of gyration corresponding to the MD optimization. (D) Contact maps before and after MD optimization. The description of the contact maps is as in Figure 3.

Together, these data underscore that INO80’s ability to orchestrate chromatin remodeling is deeply rooted in a finely balanced architecture where specific subunits combine stable cores with flexible, dynamic elements. This organizational principle allows the complex to engage nucleosomes with precision, harness ATP hydrolysis for mechanical work, and regulate gene expression and DNA repair—central processes in genome maintenance and cellular homeostasis.

#### 2.2.3 Structural Characterization of the ARP5 module

Figure 5 presents the predicted three-dimensional structure of the ARP5 module, a crucial subcomplex within the INO80 chromatin remodeling complex, composed of two subunits: Arp5 and Ies6 (Figure 5A). Our structural analysis reveals the spatial organization of Arp5 and Ies6, emphasizing regions of intrinsic flexibility enriched in residues that mediate key protein-protein interactions (Figures 5A and 5B). These interactions are further supported by crosslinking mass spectrometry data, validating their role in stabilizing the interaction interfaces within the complex.

**FIGURE 5.**
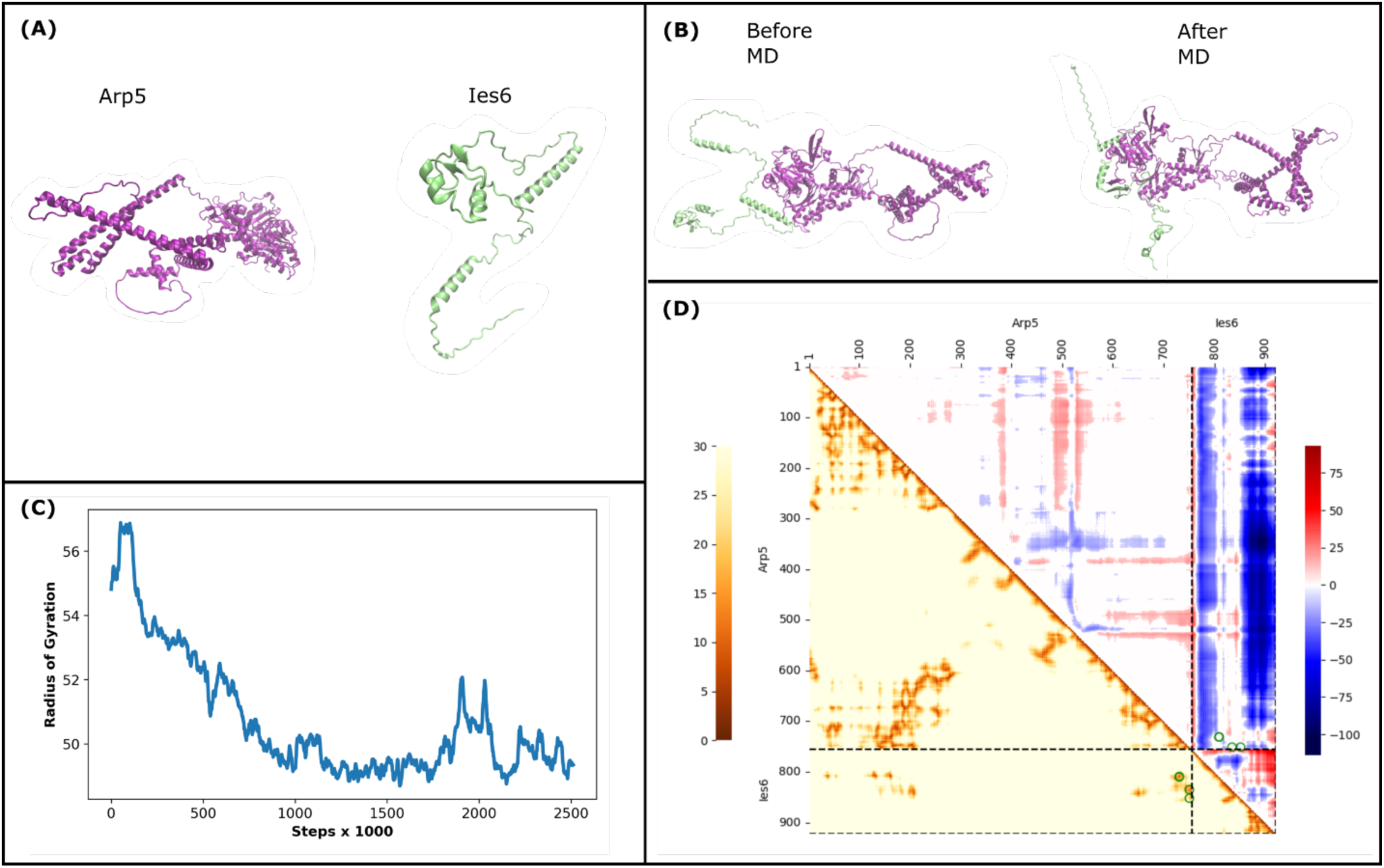
Workflow for Determining the ARP5 Module Structure: (A) Structures of individual subunits, Arp5 and Ies6, of the ARP5 subcomplex are retrieved from AlphaFold2. (B) The individual structures are assembled to form the initial template using molecular docking. (C) The radius of gyration shows the MD optimization step. (D) Contact maps before and after MD optimization. The description of the contact maps is as in Figure 3.

To capture the dynamic nature of the ARP5 module, we performed molecular dynamics simulations, enabling the initial docked model to evolve toward a conformation likely representing its physiological state (Figure 5B). This refinement led to an increased number of inter-subunit contacts and a marked reduction in the radius of gyration (Figures 5C and 5D), signifying a more compact and stable assembly. Crucially, many of the newly observed contacts in the refined structure were independently confirmed by experimental crosslinking, underscoring the synergy between computational simulations and biochemical data in resolving the module’s accurate architecture.

The novelty of our work lies in unveiling how the ARP5 module employs a flexible grappler domain to mediate nucleosome engagement, regulating INO80’s motor ATPase activity. Our findings highlight a previously uncharacterized conformational plasticity in the Arp5 subunit that facilitates differential interactions with the nucleosomal DNA entry site and histone surfaces. This dual binding mode suggests a refined mechanochemical coupling mechanism, whereby the ARP5 module not only anchors INO80 to nucleosomes but also senses DNA structural features, coordinating remodeling activity with DNA dynamics. By integrating molecular simulations with cutting-edge experimental data, this study provides unprecedented structural and mechanistic insights into the ARP5 module’s role within the INO80 complex, advancing our understanding of chromatin remodeling regulation in eukaryotic cells.

#### 2.2.4 Structural Characterization of the Ino80–Ies2 Complex

Figure 6 presents the detailed three-dimensional structure of the complex formed by the scaffolding proteins Ino80 and Ies2 (Figure 6A). The structural flexibility observed in Ino80 and Ies2, especially within loop regions, reflects the complex’s need to adapt to varying chromatin substrates and execute precise remodeling activities. Structural visualization reveals a dynamic interface where intrinsically flexible regions of both Ino80 and Ies2 mediate critical protein-protein interactions, as highlighted in Figures 6A and 6B. Within these flexible loops, several key residues were identified (Figure 6D) that are likely essential for stable association. Notably, these residues correspond closely with sites validated by independent crosslinking mass spectrometry experiments, providing strong orthogonal support for their functional roles.

**FIGURE 6.**
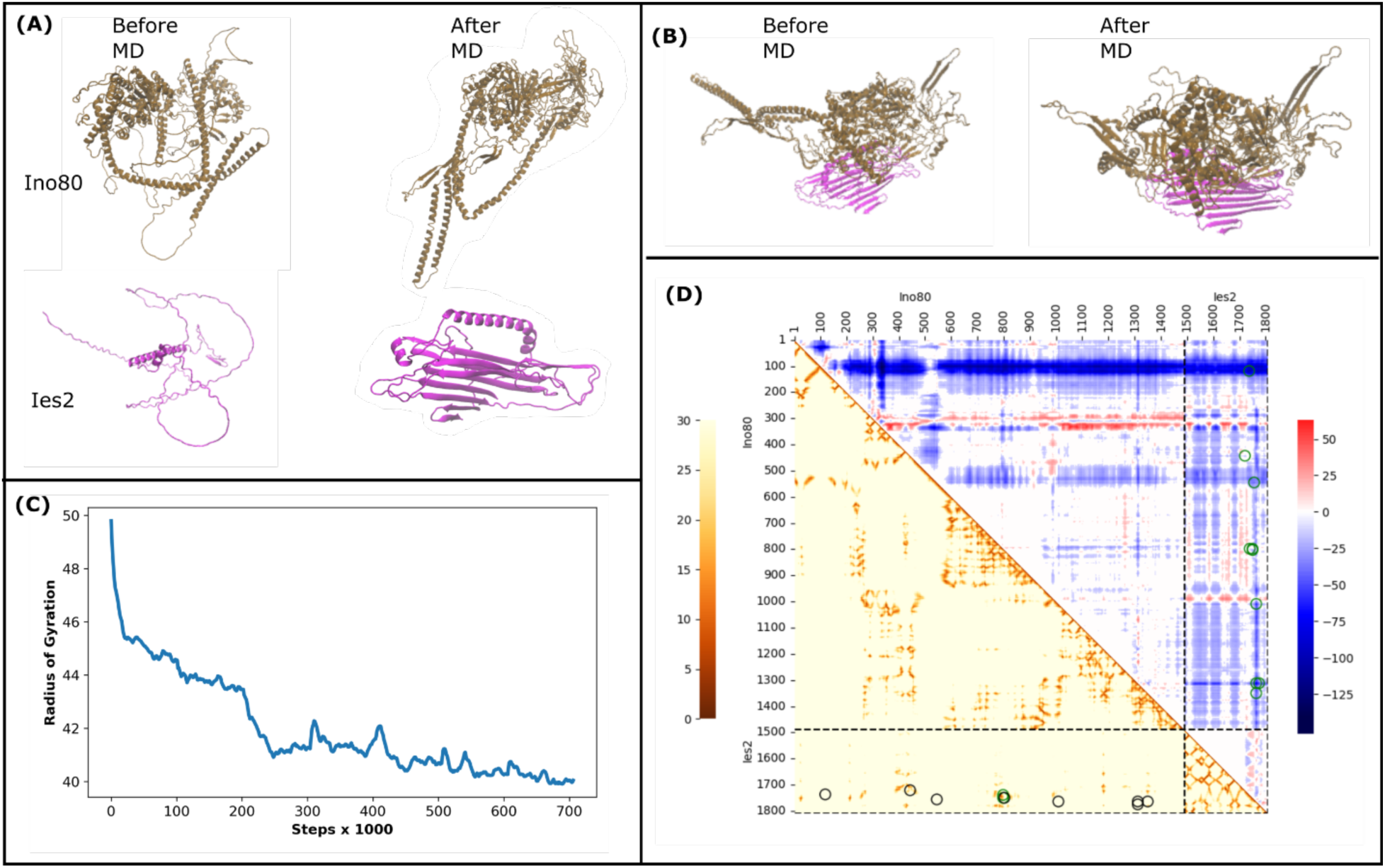
Workflow for determining the 3D structure of Ino80/Ies2 Module. (A) Structures of the individual subunits, Ino80 and Ies2, are retrieved from AlphaFold2. (B) The individual structures are assembled to form the initial template using molecular docking. (C) The radius of gyration plot corresponding to the MD optimization. (D) Contact maps before and after MD optimization. The description of the contact maps is as in Figure 3.

To refine the initial complex model, we employed extensive molecular dynamics simulations, which facilitated conformational relaxation toward a more native-like assembly (Figure 6B). This refinement revealed several hallmarks of complex stabilization: newly formed β-sheet elements emerged, suggesting an induced-fit mechanism enhancing structural rigidity (Figure 6B); the number of inter-subunit contacts increased substantially, indicative of tighter packing and improved surface complementarity (Figure 6D); and a significant decrease in the radius of gyration was observed, reflecting enhanced compaction (Figure 6C).

The novelty of this work lies in revealing that the flexible loops of Ino80 and Ies2 not only mediate binding but also undergo conformational rearrangements to adopt stabilized secondary structural elements, a process previously uncharacterized in this context. These dynamic structural adaptations likely underpin regulatory control of the INO80 complex’s ATPase activity, as the disordered domains are positioned near the motor domain, pointing to potential allosteric mechanisms.

#### 2.2.5 Structural Characterization of the RVB1/2 module

Figure 7 depicts the three-dimensional structure of the RVB1/2 module, a highly conserved and essential subcomplex of the INO80 chromatin remodeling complex, composed of the two ATPase subunits Rvb1 and Rvb2 (Figure 7A). Unlike other INO80 modules, the RVB1/2 module features a well-established, symmetric architecture, forming an alternating hexameric ring arrangement comprised of three copies each of Rvb1 and Rvb2 (Figures 7A–C). This hexameric ring structure underpins the remarkable structural stability of the module. It supports its critical role as a molecular scaffold within the INO80 complex and in multiple other chromatin remodeling assemblies. The resolved 3D structure not only confirms the evolutionary conservation of this module but offers detailed mechanistic insight into how the Rvb1/2 hexamer orchestrates the spatial organization and functional integration of INO80 subunits during nucleosome remodeling. Novel findings from our integrative model reveal the dynamic nature of Rvb1/2 subunit interfaces and demonstrate how the ATPase domain of Ino80 inserts into the central cavity of the hexamer, adopting a wheel-like configuration that mediates multivalent contacts with client subunits such as Arp5-Ies6 and Ies2. This arrangement suggests a chaperone-like function for Rvb1/2, facilitating the assembly and stability of the larger INO80 complex while potentially regulating ATPase activity through nucleotide-dependent conformational switching. Our model thereby proposes a unified mechanism linking the structural integrity of this conserved hexamer to the coordination of nucleosome engagement, ATP hydrolysis, and chromatin remodeling activity—a critical advance in understanding the multifunctionality of these AAA+ ATPases within chromatin remodelers.

**FIGURE 7.**
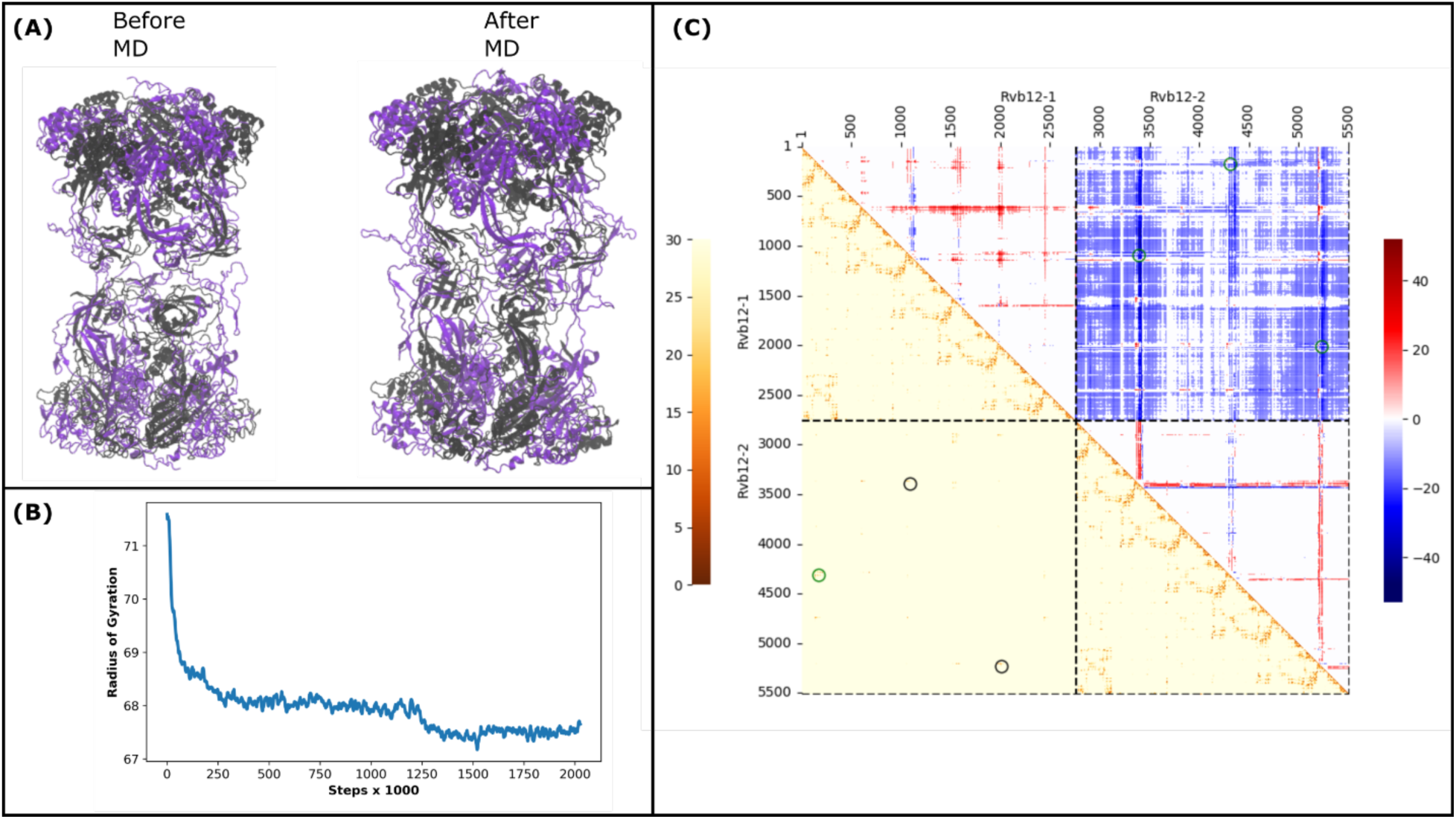
Workflow for Determining the RVB12 Dimer Structure: (A) Left: Structure after docking two Rvb1/2 monomers using HADDOCK; Right: Structure after MD optimization. (B) Radius of gyration plot corresponding to the MD optimization. (C) Contact maps before and after optimization. The description of the contact maps is as in Figure 3.

#### 2.2.6 Structural Characterization of the full-length INO80 complex

Figure 8 presents the three-dimensional structure of the full INO80 chromatin remodeling complex, assembled after comprehensive structural characterization of its four principal modules: ARP8 (Figure 5), ARP5 (Figure 6), NHP10 (Figure 4), and RVB1/2 (Figure 8). Each module represents a functionally distinct subcomplex within INO80, and these were computationally integrated into an initial full-complex template using molecular docking guided by experimentally derived crosslinking constraints. The initial assembled model was further refined through extensive molecular dynamics simulations, enabling the system to relax and extensively sample conformational space. This approach refined the inter-subunit interfaces and significantly enhanced the overall structural stability of the complex (Figure 8A). The simulations revealed the inherent flexibility and dynamic character of the complex, especially in highly mobile regions such as the interaction interface between the C-terminal domain of the Ino80 scaffold and the NHP10 module—crucial sites for mediating protein-protein interactions. The MD-refined structure displayed a notable increase in inter-subunit contacts and a decrease in radius of gyration, hallmarks of a more compact and native-like assembly (Figure 8).

**FIGURE 8.**
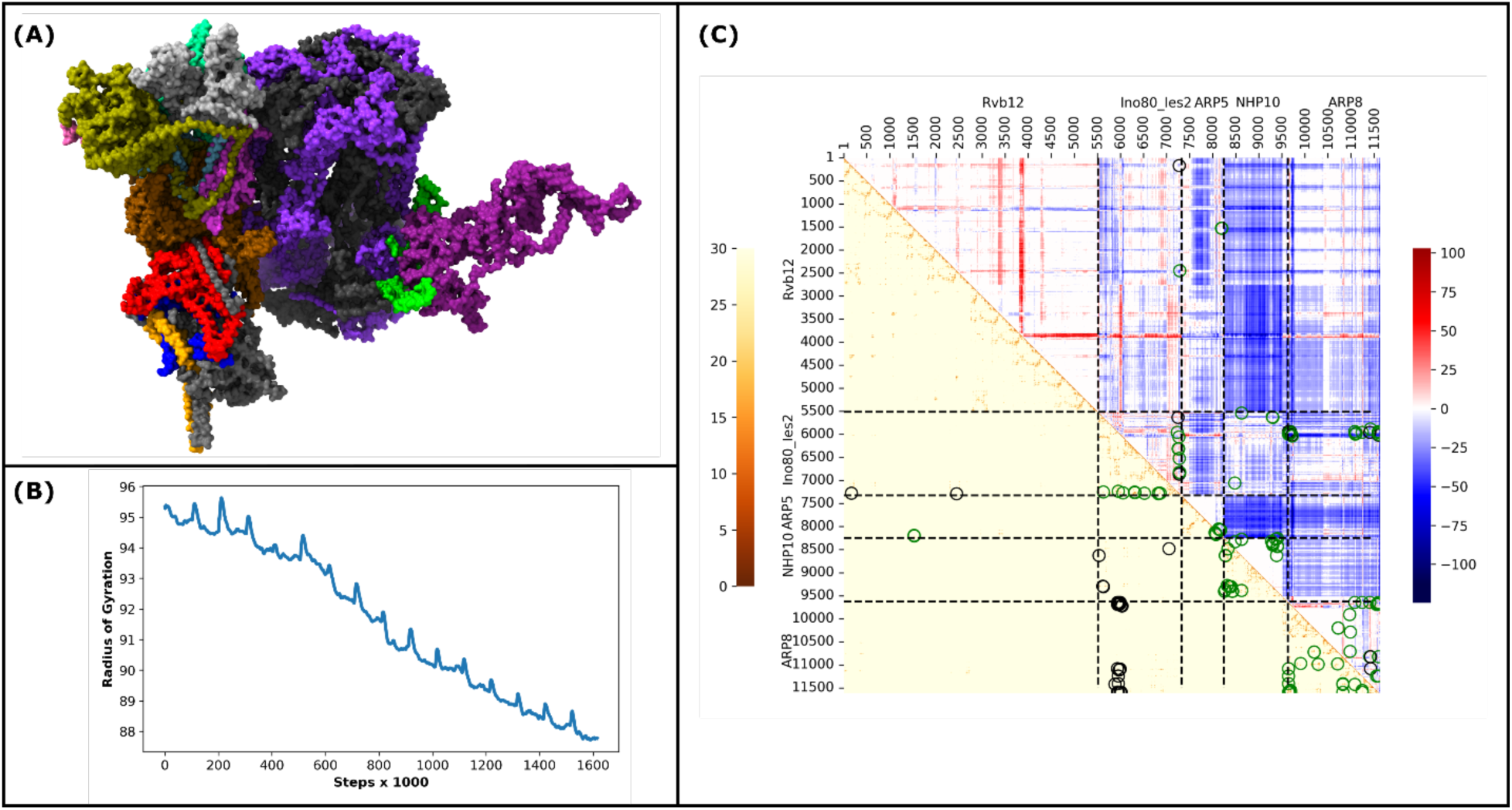
(A) MD optimized 1NO80. In the structural representation, the NHP10 module is depicted with Nhp10 in blue, Ies3 in red, Ies1 in grey, and Ies5 in orange. The ARP8 module is shown with Arp8 in tan, actin in silver, Arp4 in green, Taf14 in pink, and Ies4 in cyan. The ARP5 module includes Arp5 in purple and Ies6 in lime. Core components and additional subunits are color-coded as follows: Ino80 in ochre, Ies2 in magenta, Rvb1 in black, and Rvb2 in violet. (B) The radius of gyration plot corresponding to the MD optimization. (C) Contact maps before and after optimization. The description of the contact maps is as in Figure 3.

Integration of independent experimental crosslinking data onto the final model revealed excellent concordance between spatial proximities predicted by the structural model and observed crosslinks, thereby validating the accuracy of subunit positioning within the complex. Each module preserved its characteristic structural features while adopting an overall arrangement consistent with experimental interaction data, highlighting the modular yet interconnected architecture of INO80.

These results provide a detailed blueprint for how the structural organization of the INO80 complex underpins its diverse regulatory functions. Panel (A) demonstrates that the spatially distinctive subunit modules—for example, the hexameric Rvb1/2 core, the ATPase domains, and the flexible ARP and NHP10 modules—assemble into a defined, yet dynamic scaffold that can respond to various chromatin configurations. The contact map in panel (B) further reveals discrete intra-module interactions ensuring module integrity, while defined inter-module contacts suggest specific communication pathways across the complex. The relative separation and modularity identified here are likely critical for accommodating nucleosomes of different states and for enabling allosteric regulation, whereby DNA engagement at one domain can trigger conformational changes and enzymatic activation at another. This architecture enables the INO80 complex to reposition, slide, or evict nucleosomes, facilitating DNA repair or transcriptional control in response to cellular cues. Thus, our integrative modeling delineates not just a static structure but a functional platform that reconciles the requirements for biochemical precision, substrate diversity, and the regulatory adaptability expected of a central chromatin remodeler in genome regulation.

This work represents a significant advancement, as it provides the first high-resolution integrative model capturing both the spatial organization and dynamic behavior of the entire INO80 complex. Our structural elucidation of the elusive NHP10 module, and its dynamic tethering via the extended N-terminal tail of Ino80, sheds light on regulatory mechanisms potentially critical for function. Additionally, the refined full-length model illuminates mechanistic insights into how modular flexibility and structural cohesion enable chromatin remodeling activities fundamental to gene regulation, DNA repair, and replication. Importantly, our integrative approach—combining crosslinking mass spectrometry, AI-driven structural prediction (AlphaFold), molecular docking, and molecular dynamics—circumvents limitations of classical structural techniques that struggle with flexible, dynamic protein assemblies. This framework is broadly applicable to other large, multi-subunit complexes and sets the stage for future studies dissecting the mechanistic roles of chromatin remodelers in health and disease.

In summary, this study not only advances molecular understanding of INO80 complex but exemplifies the power of hybrid integrative structural biology methods in revealing the architecture and function of dynamic macromolecular machines, forging new paths toward therapeutic targeting of chromatin-related disorders.

## 3. DISCUSSION

Our integrative structural analysis of the INO80 chromatin remodeling complex provides a comprehensive three-dimensional framework that elucidates both the spatial organization and dynamic interplay of its principal modules: ARP8, ARP5, NHP10, and RVB1/2. By synergistically combining experimental crosslinking mass spectrometry, state-of-the-art AI-based structural predictions from AlphaFold, molecular docking, and rigorous molecular dynamics simulations, we have generated a high-resolution model that aligns closely with empirical data and bridges critical knowledge gaps left unresolved by conventional structural biology methods.

Traditional techniques such as X-ray crystallography and cryo-EM have been invaluable for resolving the stable core of INO80, such as the RUVBL1/2 AAA+ hexamer and the Swi2/Snf2 ATPase motor (14). However, they have historically struggled to define highly flexible and heterogeneous regions, particularly the elusive NHP10 module and intrinsically disordered scaffold extensions. Our integrative approach leverages the atomic precision of AI-driven predictions, the spatial anchoring power of crosslinking restraints, and the dynamic sampling capabilities of molecular dynamics simulations to resolve these flexible regions, capturing the conformational plasticity essential for INO80’s biological function.

For example, the flexible tethering of the NHP10 subcomplex to the extended N-terminal tail of the Ino80 scaffold appears critical for regulatory subunit crosstalk and allosteric modulation of ATPase activity. Our refined model also reveals the dynamic spatial relationships of the ARP8 and ARP5 modules, shedding light on their roles as DNA sensors and mechanochemical transducers that govern nucleosome engagement and remodeling. Meanwhile, the RVB1/2 hexamer emerges as a central scaffolding hub, mediating multivalent interactions that stabilize the complex and couple ATP hydrolysis to nucleosome sliding.

These mechanistic insights go beyond earlier static or partial structural characterizations, demonstrating how modular flexibility, dynamic interfaces, and finely tuned allosteric regulation converge to enable the multifaceted functions of INO80 in chromatin remodeling, DNA damage repair, and transcription regulation. Notably, our model complements recent high-resolution cryo-EM structures of the human INO80 core, extending understanding by resolving flexible and previously uncharacterized modules, particularly the NHP10 unit and its intricate tethering mechanisms.

Comparative analysis with related remodelers such as RSC and SWR-C highlights INO80’s distinct architectural and functional adaptations (32). Unlike SWR-C’s more rigid substrate specificity for ATP-dependent deposition of the H2A.Z variant, INO80 is uniquely versatile, capable of both nucleosome mobilization and histone variant dimer exchange. This dual functionality is underpinned by the pronounced conformational flexibility of subcomplexes like NHP10 and the ARP modules, which facilitate regulated allosteric communication and adaptive nucleosome interactions. Our findings underscore evolutionary divergence among remodeler families, emphasizing the role of dynamic modularity in tailoring remodeling activities to specific genomic and cellular contexts.

More broadly, our integrative workflow establishes a versatile paradigm for structural biology, effectively complementing and extending existing computational and experimental approaches. This robust integrative modeling workflow has been encapsulated into a versatile software package named INTEGRATOR (INTEGRAtive TempOral and stRuctural Analysis of pROtein modules), designed to facilitate the structural elucidation of large, dynamic macromolecular assemblies. INTEGRATOR is adaptable to a broad range of large, flexible protein complexes and chromatin-associated assemblies whose dynamic and heterogeneous nature challenges traditional single-method structural approaches, offering a comprehensive platform for integrative, multi-modal structural biology. As the field advances toward elucidating entire molecular machines in their functional states, integrative methods such as ours will be indispensable for dissecting the molecular basis of genome regulation and its dysregulation in disease.

## 4 CONCLUSION

Our integrative structural analysis advances the understanding of the INO80 chromatin remodeling complex by revealing how its diverse modules cooperate in a dynamically orchestrated molecular machine. By harnessing crosslinking mass spectrometry, AI-driven structure prediction, molecular docking, and molecular dynamics within the INTEGRATOR software platform, we deliver a unified model that resolves previously inaccessible regions while capturing essential conformational flexibility. Importantly, the adaptability of INTEGRATOR makes it possible to tackle a wide array of dynamic, heterogeneous protein assemblies across chromatin biology and other fields where molecular plasticity is central to biological regulation. With this platform, structural biologists can move beyond static snapshots to holistic, physiologically relevant models that couple spatial and temporal resolution. Looking ahead, our framework encourages integration of new data types and methodological approaches, setting the stage for deeper mechanistic insight into genome regulation and molecular disease processes.

## 5. MATERIALS AND METHODS

### 5.1 Protein Structure Prediction Workflow

Our study introduces a robust, multi-stage integrative modeling workflow designed to elucidate the architecture of large, flexible, and multi-subunit protein assemblies that are challenging for traditional structural methods. The framework begins with the generation of initial structural models for individual subunits, primarily leveraging predictions from AlphaFold2 (24). These subunits are then assembled into functional subcomplexes using a sequential, pairwise docking strategy implemented in HADDOCK (31). This assembly is not arbitrary; it is guided by a combination of HADDOCK’s scoring function and, where available, experimentally derived distance restraints from crosslinking mass spectrometry (XL-MS), ensuring that the initial models are physically plausible and consistent with empirical data. In the absence of cross-linking data, we can leverage a topological scoring approach to infer probable direct interactions (9, 33).

The second stage of our workflow involves extensive structural refinement and optimization through molecular dynamics (MD) simulations (34). Each subcomplex is first subjected to an unrestrained MD simulation where structural memory restraints, weighted by AlphaFold confidence scores, guide the system toward a stable low-energy conformation. Following this initial relaxation, we progressively enforce experimental XL-MS distance restraints in a stepwise manner. This iterative process systematically refines the subcomplex architecture, correcting inter-atomic distances to satisfy the crosslinking data and significantly improving the model’s accuracy.

The final stage consists of assembling the refined subcomplexes into the complete macromolecular machine, again using a crosslink-guided docking approach, followed by a final round of iterative MD optimization on the entire complex. This hierarchical strategy, moving from individual subunits to subcomplexes and finally to the whole assembly, effectively navigates the vast conformational space. The final model’s validity is rigorously assessed by its high concordance with the independent XL-MS dataset and the convergence of structural stability metrics like the radius of gyration. This integrative paradigm provides a generalizable and powerful framework for determining high-quality structural models of dynamic and heterogeneous molecular machines.

### 5.2 Subcomplex preparation

The Rvb1/2 monomer structure was initially obtained from the Protein Data Bank (PDB ID: 8ETW). To complete missing residues, PyMol (35) was employed to model and incorporate absent regions. The modified protein structure was then subjected to optimization using Associative memory, Water-mediated, Structure and Energy Model Molecular Dynamics (AWSEM-MD) (36) for 100,000 simulation steps. A critical parameter influencing the MD optimization is the spring constant applied as a restraining potential, which enforces structural memory restraint during simulation. To identify the optimal spring constant, we conducted a systematic series of AWSEM-MD simulations on the modified Rvb1/2 monomer, varying the restraint force constant across values of 1, 10, 20, 30, 40, 50, 60, 70, 80, 90, and 100 kcal/mol/Å². The associative memory term was applied to all residues except those introduced through PyMol remodeling. Post-optimization, the root mean square deviation (RMSD) of atomic coordinates was computed to assess structural convergence and stability. The results, summarized in Supplementary Figure 1, display how increasing restraint strength affects the structural deviation, allowing us to select an optimal parameter that balances structural fidelity and conformational flexibility. Based on the results shown in Supplementary Figure 1, we selected a spring constant (K) value of 30 kcal/mol/Å² for our molecular dynamics (MD) optimization, as this restraint strength yielded sufficiently small deviations in residue positions that we deem most reliable (Supplementary Figure 2). To construct the Rvb1/2 dimer structure, the optimized monomer model was duplicated, and the two monomers were docked together using HADDOCK 3.0. The docking was guided by three experimentally validated cross-links (Table 1), which specifically connect the bottom regions of one Rvb1/2 monomer to the other.

**Table 1:** Cross-links used to assemble the Rvb1/2 dimer.

| Dimer 1<br>(residue number) | Dimer 2<br>(residue number) |
| --- | --- |
| 177 | 1558 |
| 2015 | 2477 |
| 1096 | 639 |

HADDOCK 3.0 employs a multi-stage docking workflow, consisting of rigid body docking, flexible refinement, and clustering steps. For this study, we utilized the rigid body docking stage results to balance computational efficiency with reasonable structural estimates for subsequent MD simulations. To identify the optimal docked conformation, we evaluated the average distances between the Cα atoms corresponding to the three cross-link restraints connecting the two monomers. We selected the structure showing the minimum average distance. This approach ensured adherence to experimental constraints while providing a robust starting point for dynamic refinement.

#### 5.2.1 NHP10

The NHP10 subcomplex is composed of four protein subunits: Nhp10, Ies3, Ies1, and Ies5. Initial structural models of these individual proteins were obtained from AlphaFold. To assemble the NHP10 complex, we employed HADDOCK 3.0, utilizing a sequential docking strategy in which two subunits were docked at a time. The docking order proceeded as follows: first, Nhp10 was docked with Ies3; next, the resulting Nhp10-Ies3 complex was docked with Ies1; finally, the Nhp10-Ies3-Ies1 assembly was docked with Ies5. For each docking step, 100 initial rigid-body docking configurations were generated and evaluated. The highest-ranked structure at the final stage of HADDOCK docking, as determined by HADDOCK’s scoring function, was selected as the representative model for that step and carried forward to the next docking stage. This approach enabled stepwise construction of a stable NHP10 complex conformation consistent with predicted interaction interfaces and experimental data.

#### 5.2.2 ARP8

The ARP8 subcomplex consists of five protein subunits: Arp8, Act, Arp4, Taf14, and Ies4. Structural models for each subunit were obtained from AlphaFold. Assembly of the ARP8 complex was performed using a sequential docking strategy analogous to that applied for the NHP10 complex. Docking proceeded in the following order: first, Arp8 was docked with Act; next, the resulting Arp8-Act complex was docked with Arp4; this was followed by docking of Taf14 to the Arp8-Act-Arp4 assembly; finally, Ies4 was docked to the fully assembled Arp8-Act-Arp4-Taf14 complex. At each docking step, 100 initial rigid-body configurations were generated and evaluated. The highest-ranking model at the final stage of each HADDOCK docking cycle, as determined by HADDOCK’s scoring function, was selected as the representative structure and carried forward. This stepwise approach enabled systematic construction of a stable, physically plausible ARP8 complex architecture consistent with predicted interaction interfaces.

#### 5.2.3 ARP5

The ARP5 subcomplex consists of two proteins, Arp5 and Ies6. Structural models of these subunits were obtained from AlphaFold. Unlike the other complexes, the ARP5 assembly required only a single docking step. During this step, 100 initial rigid-body docking configurations were generated and evaluated using HADDOCK 3.0. The highest-ranked structure at the final docking stage, as determined by HADDOCK’s scoring criteria, was selected as the representative structure for the ARP5 complex. This streamlined docking approach efficiently produced a reliable model suitable for subsequent molecular dynamics refinement.

#### 5.2.4 Ino80+Ies2

Initially, we considered attaching the Ino80 and Ies2 subunits independently to the other INO80 subcomplexes. However, the large number of cross-links between Ino80 and the Rvb1/2 module made identifying a precise attachment site challenging. This complexity was compounded by the fact that each residue index in Rvb1/2 corresponds to three separate protein chains (Rvb1 is represented by chains A, C, and E; Rvb2 by chains B, D, and F), complicating the assignment of interaction interfaces. In contrast, the cross-links connecting Ies2 to Rvb1/2 were fewer in number and more specific, facilitating easier interpretation. Leveraging this, we adopted a stepwise assembly strategy: first, Ino80 was docked and connected to Ies2; next, the Ino80+Ies2 subcomplex was attached to the Rvb1/2 module, focusing exclusively on the two unambiguous cross-links between Ies2 and Rvb1/2. These two cross-links, detailed in Table 2, served as robust spatial restraints for this final docking step, facilitating accurate anchoring of the Ino80+Ies2 subcomplex onto the Rvb1/2 hexameric scaffold.

**Table 2:**
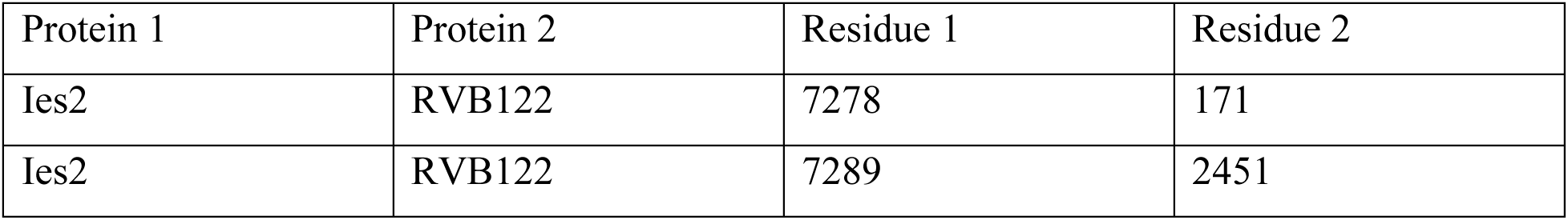
Cross-links considered for (Ino80+Ies2)-Rvb1/2 docking.

### 5.3 MD optimization

#### 5.3.1 Optimizing the subcomplexes

Molecular dynamics (MD) optimization using AWSEM was performed for each subcomplex, including Ino80 and Ies2, over 2 million simulation steps without applying any restraints from mass spectrometry (MS) cross-links. During this phase, residues with AlphaFold confidence scores above 70% were incorporated into the memory file to guide the structural sampling. Subsequently, an additional 200,000 MD steps were executed, during which only residues with confidence scores exceeding 90% were included in the memory file, allowing focused refinement of high-confidence regions. To facilitate docking, the Ino80 and Ies2 proteins were combined into a single subcomplex, which was then docked to the RVB1/2 module using 100 initial rigid-body configurations. This step yielded five subcomplexes in total: RVB1/2, NHP10, ARP8, ARP5, and Ino80+Ies2. Further optimization of these subcomplexes was conducted by progressively enforcing MS cross-link-derived distance restraints in a stepwise manner. If the Cα–Cα distance of a given cross-link was less than 10 Å, no additional restraint was applied. For distances exceeding 10Å, a harmonic restraint was enforced, setting the equilibrium distance to 70% of the initial HADDOCK-predicted distance. If this calculated equilibrium distance was below 10Å, it was set to 9.5 Å to ensure a physically reasonable restraint. This iterative process continued until all cross-link distances were reduced below 10Å. In cases where the initial HADDOCK distance was under 10Å but subsequently increased beyond this threshold due to the lack of restraints, the equilibrium distance was set to 9.5 Å to correct this deviation.

#### 5.3.2 Optimizing INO80

The INO80 complex was assembled through a sequential docking strategy involving the RVB1/2 dimer, Ino80-Ies2, ARP5, NHP10, and ARP8 subcomplexes. Initially, the RVB1/2 dimer was docked to the Ino80-Ies2 subcomplex, followed by docking the resulting assembly with the ARP5 subcomplex. This intermediate complex was then docked with NHP10, and finally, the entire assembly was combined with the ARP8 subcomplex. During iterative optimization, the number of satisfied mass spectrometry (MS) cross-links was monitored to assess structural improvements relative to experimental data. The results, shown in Supplementary Figure 3, demonstrate that molecular dynamics relaxation combined with enforced MS cross-link restraints significantly increased the proportion of cross-links with Cα–Cα distances under 26Å, a critical threshold corresponding to the typical maximal span of the disuccinimidyl suberate (DSS) crosslinker(37, 38). Specifically, our refined model satisfied over 90% of the cross-links below 24Å, a substantial improvement compared to the initial HADDOCK model, which satisfied only about 50% of cross-links at this threshold. These findings highlight the effectiveness of our integrative modeling approach in producing a native-like, experimentally consistent model of the full INO80 complex.

**Table 3:** Cross-links considered for subcomplex and INO80 optimization.

| Protein1 | Protein2 | Res1 | Res2 |
| --- | --- | --- | --- |
| Ies2 | RVB122 | 7278 | 171 |
| Ies2 | RVB122 | 7289 | 2451 |
| Ies6 | RVB122 | 8192 | 1529 |
| Ies6 | RVB122 | 8204 | 1532 |
| Ies2 | Ino80 | 7235 | 5957 |
| Ies2 | Ino80 | 7251 | 5633 |
| Ies2 | Ino80 | 7253 | 6312 |
| Ies2 | Ino80 | 7264 | 6312 |
| Ies2 | Ino80 | 7264 | 6317 |
| Ies2 | Ino80 | 7270 | 6059 |
| Ies2 | Ino80 | 7278 | 6523 |
| Ies2 | Ino80 | 7278 | 6826 |
| Ies2 | Ino80 | 7278 | 6864 |
| Ies2 | Ino80 | 7289 | 6826 |
| Ies3 | Ino80 | 8629 | 5538 |
| Ies1 | Ino80 | 9295 | 5622 |
| Ies1 | Ino80 | 9306 | 5633 |
| Arp8 | Ino80 | 9646 | 5941 |
| Arp8 | Ino80 | 9646 | 5974 |
| Arp8 | Ino80 | 9646 | 5997 |
| Arp8 | Ino80 | 9672 | 5957 |
| Arp8 | Ino80 | 9672 | 5974 |
| Arp8 | Ino80 | 9679 | 5957 |
| Arp8 | Ino80 | 9679 | 5992 |
| Arp8 | Ino80 | 9690 | 5941 |
| Arp8 | Ino80 | 9690 | 5957 |
| Arp8 | Ino80 | 9690 | 5970 |
| Arp8 | Ino80 | 9690 | 5974 |
| Arp8 | Ino80 | 9716 | 6019 |
| Arp8 | Ino80 | 9725 | 6037 |
| Arp4 | Ino80 | 11083 | 5938 |
| Arp4 | Ino80 | 11093 | 5992 |
| Arp4 | Ino80 | 11093 | 5997 |
| Arp4 | Ino80 | 11113 | 5974 |
| Arp4 | Ino80 | 11243 | 5957 |
| Ino80 | Taf14 | 5884 | 11420 |
| Ino80 | Taf14 | 5957 | 11408 |
| Ies4 | Ino80 | 11592 | 5941 |
| Ies4 | Ino80 | 11596 | 5957 |
| Ies4 | Ino80 | 11596 | 6019 |
| Ies4 | Ino80 | 11597 | 5941 |
| Ies4 | Ino80 | 11597 | 5957 |
| Ies4 | Ino80 | 11597 | 5997 |
| Ies4 | Ino80 | 11597 | 6001 |
| Ies2 | Ies3 | 7056 | 8481 |
| Arp5 | Ies6 | 8054 | 8132 |
| Arp5 | Ies6 | 8054 | 8133 |
| Arp5 | Ies6 | 8074 | 8158 |
| Arp5 | Ies6 | 8074 | 8175 |
| Ies3 | Nhp10 | 8486 | 8331 |
| Ies3 | Nhp10 | 8629 | 8276 |
| Ies3 | Nhp10 | 8629 | 8279 |
| Ies1 | Nhp10 | 9277 | 8318 |
| Ies1 | Nhp10 | 9306 | 8353 |
| Ies1 | Nhp10 | 9306 | 8381 |
| Ies1 | Nhp10 | 9306 | 8404 |
| Ies1 | Nhp10 | 9371 | 8276 |
| Ies5 | Nhp10 | 9405 | 8258 |
| Ies5 | Nhp10 | 9405 | 8422 |
| Ies5 | Nhp10 | 9405 | 8436 |
| Ies1 | Ies3 | 9387 | 8629 |
| Act | Arp8 | 10721 | 10202 |
| Act | Arp8 | 10723 | 10202 |
| Arp4 | Arp8 | 10971 | 9910 |
| Arp4 | Arp8 | 10988 | 10289 |
| Arp4 | Arp8 | 11083 | 9646 |
| Arp4 | Arp8 | 11243 | 9646 |
| Arp8 | Taf14 | 9646 | 11408 |
| Arp8 | Ies4 | 9646 | 11597 |
| Arp8 | Ies4 | 9672 | 11597 |
| Arp8 | Ies4 | 9679 | 11596 |
| Arp8 | Ies4 | 9679 | 11597 |
| Arp8 | Ies4 | 9686 | 11596 |
| Arp8 | Ies4 | 9686 | 11597 |
| Arp8 | Ies4 | 9690 | 11554 |
| Arp8 | Ies4 | 9690 | 11560 |
| Arp8 | Ies4 | 9690 | 11587 |
| Arp8 | Ies4 | 9690 | 11596 |
| Arp8 | Ies4 | 9690 | 11597 |
| Act | Arp4 | 10710 | 10980 |
| Arp4 | Taf14 | 10824 | 11408 |
| Arp4 | Taf14 | 10824 | 11425 |
| Arp4 | Taf14 | 11083 | 11425 |
| Arp4 | Ies4 | 10824 | 11597 |
| Arp4 | Ies4 | 11243 | 11554 |
| Arp4 | Ies4 | 11243 | 11592 |
| Arp4 | Ies4 | 11243 | 11596 |
| Arp4 | Ies4 | 11243 | 11597 |

### 5.4 Model Validation

Validating the accuracy of final protein structures following molecular dynamics (MD) simulations remains a critical step, particularly for large, flexible complexes like INO80, where high-resolution reference structures are unavailable. Traditional validation approaches often rely on direct comparison to an experimentally determined initial structure or the fitting of models into cryo-EM or cryo-ET density maps. However, these methods become impractical for highly dynamic systems such as INO80 due to their intrinsic conformational heterogeneity.

To rigorously validate our final INO80 model, we employed a multi-faceted strategy. First, we compared contact maps derived from MD simulation trajectories, which reflect residue-residue connectivity and module interactions, against independent crosslinking mass spectrometry (XL-MS) data. This comparison provides a robust, experimental benchmark for assessing the fidelity of inter-subunit contacts and overall architectural consistency.

In parallel, we monitored the radius of gyration throughout the simulations, observing its stabilization to a near-constant value, a hallmark of system convergence and structural stability. This combination of spatial connectivity validation and dynamic stability assessment offers compelling evidence that our integrative modeling approach produces a biologically relevant and stable representation of the INO80 assembly.

### 5.5 Model Visualization

3D structures were visualized using the Visual Molecular Dynamics (VMD) software package (39) or Chimera software (40). Contact maps and other plots were generated using the Python scripts, which are available at this GitHub link.

---

**FIGURE 9.** The workflow for predicting the INO80 structure involves several steps. First, the protein PDBs are obtained. If they are not available in the Protein Data Bank, they are retrieved from AlphaFold. Next, the complexes are created using HADDOCK, and MD optimization is performed on each sub-complex. Subsequently, HADDOCK is used to assemble the INO80 complex. Finally, MD optimization is conducted on the INO80 complex.

**FIGURE 10.** Workflow of the integrative modeling of the INO80 complex. (A) PDBs of the individual subunit of the INO80 complex (B). Shared and non-shared subunits of the yeast and human INO80 complex; (C) Individual modules of the INO80 complex (C) are assembled into the final complex (D). In the structural representation, the NHP10 module is depicted with Nhp10 in blue, Ies3 in red, Ies1 in grey, and Ies5 in orange. The ARP8 module is shown with Arp8 in tan, actin in silver, Arp4 in green, Taf14 in pink, and Ies4 in cyan. The ARP5 module includes Arp5 in purple and Ies6 in lime. Core components and additional subunits are color-coded as follows: Ino80 in ochre, Ies2 in magenta, Rvb1 in black, and Rvb2 in violet. This color scheme highlights the modular architecture of the INO80 complex and facilitates visual distinction between conserved modules and subunits across structural figures.

**FIGURE 11.** Workflow for Determining the NHP10 Module Structure: (A) Structures of the subunits of the ARP8 subcomplex are retrieved from AlphaFold2. (B) The individual structures are assembled to form the initial template using molecular docking. (C) The radius of gyration plot corresponding to the MD optimization. (D) Contact maps before and after MD optimization. The lower triangle shows the residue-residue distances; distances corresponding to the cross-links given in Tosi et al. (10) are marked in circles. Distances greater than 24 Å are colored in black, while others are in green. The upper triangle depicts the difference between final and initial cross-link distances.

**FIGURE 12.** Workflow for Determining the ARP8 Module Structure: (A) Structures of the individual subunits of the ARP8 subcomplex are retrieved from AlphaFold2. (B) The individual structures are assembled to form the initial template using molecular docking. (C) The radius of gyration corresponding to the MD optimization. (D) Contact maps before and after MD optimization. The description of the contact maps is as in Figure 3.

**FIGURE 13.** Workflow for Determining the RVB12 Dimer Structure: (A) Left: Structure after docking two Rvb1/2 monomers using HADDOCK; Right: Structure after MD optimization. (B) Radius of gyration plot corresponding to the MD optimization. (C) Contact maps before and after optimization. The description of the contact maps is as in Figure 3.

**FIGURE 14.** (A) MD optimized 1NO80. In the structural representation, the NHP10 module is depicted with Nhp10 in blue, Ies3 in red, Ies1 in grey, and Ies5 in orange. The ARP8 module is shown with Arp8 in tan, actin in silver, Arp4 in green, Taf14 in pink, and Ies4 in cyan. The ARP5 module includes Arp5 in purple and Ies6 in lime. Core components and additional subunits are color-coded as follows: Ino80 in ochre, Ies2 in magenta, Rvb1 in black, and Rvb2 in violet. (B) The radius of gyration plot corresponding to the MD optimization. (C) Contact maps before and after optimization. The description of the contact maps is as in Figure 3.

## AUTHOR CONTRIBUTIONS

**Jules Nde:** writing –original draft; formal analysis; data curation; validation; visualization. **Gihan Panapitiya**: writing –original draft; formal analysis; data curation; validation; visualization. **Margaret S. Cheung:** Conceptualization; methodology, writing – review and editing; computational platform administrator. **Mark Maupin**: Conceptualization; methodology; writing –original draft; investigation, supervision. **Mihaela Sardiu:** Conceptualization; methodology; writing; investigation, supervision, funding acquisition.

## ACKNOWLEDGMENTS

This research was performed on a project award (10.46936/lser.proj.2023.60699/60008913) from the Environmental Molecular Sciences Laboratory, a DOE Office of Science User Facility sponsored by the Biological and Environmental Research program under Contract No. DE-AC05-76RL01830. The authors also thank Amity Andersen’s effort in modifying the data transformation in the workflow. J.N. and M.S.C. also thanks the National Science Foundation (MCB 2221824) for partially supporting J.N’s graduate research work at the University of Washington, Seattle.

